# Mathematical Modeling of Fluconazole Resistance in the Ergosterol Pathway of *Candida albicans*

**DOI:** 10.1101/2022.06.09.495576

**Authors:** Paul K. Yu, Llewelyn S. Moron, Angelyn R. Lao

## Abstract

Candidiasis is reported as the most common fungal infection in the critical care setting. The causative agent of this infection is a commensal pathogen belonging to the genus *Candida*, most common species of which is the *Candida albicans*. The ergosterol pathway in yeast is a common target by many antifungal agents since ergosterol is an essential component of the cell membrane. The current antifungal agent of choice for the treatment of Candidiasis is fluconazole, which is classified under the azole antifungals. In recent years, the significant increase of fluconazole-resistant *C. albicans* in clinical samples calls for a need to search for other possible drug targets. In this study, we constructed a mathematical model of the ergosterol pathway of *C. albicans* using ordinary differential equations with mass action kinetics. From the model simulations, we found the following results: (1) a partial inhibition of the sterol-methyltransferase enzyme yields a fair amount of fluconazole resistance, (2) an overexpression of the *ERG6* gene, leading to increased sterol-methyltransferase enzyme, is a good target of antifungals as an adjunct to fluconazole, (3) a partial inhibition of lanosterol yields a fair amount of fluconazole resistance, (4) the C5-desaturase enzyme is not a good target of antifungals as an adjunct to fluconazole, (5) the C14α-demethylase enzyme is confirmed to be a good target of fluconazole, and (6) the dose-dependent effect of fluconazole is confirmed. This study hopes to aid experimenters narrow down the possible drug targets prior to doing costly and time-consuming experiments, and to serve as a cross-validation tool for experimental data.

## 1. Introduction

Candidiasis refers to a broad range of fungal infections caused by different species of the fungal commensal pathogen belonging to the genus *Candida*. The following *Candida* species are all reported to cause cutaneous, mucosal, bloodstream, and deep-seated tissue infections: *Candida albicans, C. glabrata, C. tropicalis, C. parapsilosis*, and *C. krusei*. The most common of which is the Candida *albicans*. [1]

Invasive candidiasis is the term used for the more concerning type that includes the bloodstream yeast infections, also known as candidemia. Likewise, deep-seated tissue infections are infections that occur in sterile sites such as the peritoneal cavity, abdominal and non abdominal sites, namely the bones, muscles, joints, eyes, or central nervous system [2]. Surprisingly, invasive type is the most common fungal infection in the critical care setting, with a crude mortality of about 40 to 55 percent [3].

Logan et al. (2020) noted several risk factors for invasive candidiasis that include the use of broad-spectrum antimicrobials, immunosuppressive drugs, total parenteral nutrition, iatrogenic interventions, along with several challenges in the critical care setting, namely, the shift to more resistant *Candida* epidemiology, and the appropriate strategies for antifungal therapy [3]. Among the different classes of antifungal agents used for invasive candidiasis are agents such as azoles, polyenes, echinocandins, and allylamines. However, fluconazole, a fungistatic drug belonging to the azole group, is currently the preferred treatment for *C. albicans* infections due to its low cost, limited toxicity, and oral administration [4]. The mechanism of action of azole drugs is the inhibition of the C14α-demethylase enzyme, thus blocking the ergosterol biosynthesis pathway, which is an essential component of the fungal cell membrane in *Candida* species [5].

In recent years, Whaley et al. (2017) noted a high-level of resistance to azoles among *Candida* species and some non-albicans *Candida* species have already reported with intrinsic resistance to azoles [4]. In fact, the Centers for Disease Control and Prevention (CDC) in its Antibiotic Resistance Threats in the United States, 2019 report classifies drug-resistant *Candida* as one of the serious threats, with an estimate of 34,800 cases and 1,700 deaths [6]. They noted that about 7% of the all *Candida* blood samples tested are resistant to fluconazole [6].

There is an abundant amount of experimental investigations in azole resistance mechanisms in *C. albicans*, which can be costly and time-consuming. In this study, we propose a different approach to investigating fluconazole resistance in *C. albicans*, using mathematical models to determine possible drug targets within the ergosterol pathway. Its aim is to narrow down possible drug targets prior to doing costly and time-consuming experiments, and to serve as a cross-validation tool for experimental data. To our knowledge, this is the first study to investigate the fluconazole resistance in the ergosterol pathway of *Candida albicans* using mathematical modeling.

The rest of the paper is organized as follows: In section 2, we discuss, in detail, the mathematical modeling process. The results of the model simulations are shown in section 3. Then, it is followed by a discussion on how the results compare with previous literature, in section 4. Lastly, section 5 draws the conclusion of the study and proposes further recommendations.

## 2. Mathematical Modeling

The model for the ergosterol biosynthesis pathway, as shown in Figure 1, in *C. albicans* is built adopting the pathways of Sanglard et al. (2003) (see Figure 1 in [5]), and Martel et al. (2010) (see Figure 1 in [7]). We noted several differences between the two studies, notably, the substrate for the C14α-demethylase enzyme is lanosterol in Sanglard et al. (2003), however, it is eburicol in the pathway presented by Martel et al. (2010). There are still disparities in the literature regarding this matter. In the study, we followed Sanglard et al. (2003) for the construction of our model.

**Figure 1.**
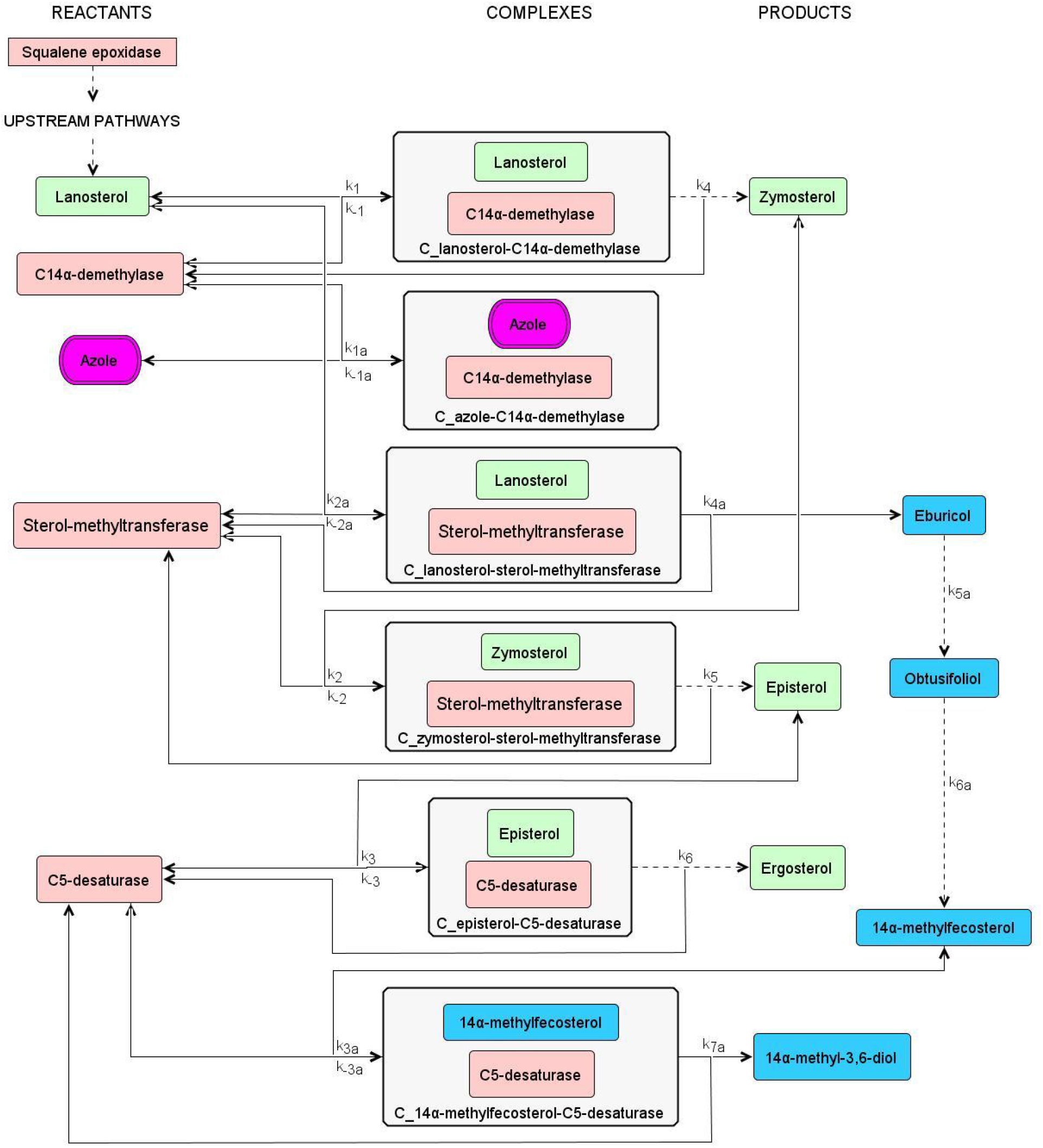
Ergosterol biosynthesis pathway. The pathway starts from lanosterol to ergosterol synthesis. It does not include the pathways upstream of lanosterol, such as the squalene epoxidase enzyme. The metabolites in green denote the ergosterol pathway in *C. albicans* when azole is absent, which leads to its survival. Whereas, those in blue represent the alternative pathway during azole inhibition, which leads to its demise due to the production of the toxic sterol, 14α-methyl-3,6-diol. The enzymes are colored in pink. The azole drug is colored in magenta. The arrows pertain to the direction of the reaction, such that bi-directional arrows indicate reversible reactions.

Due to the complexity of the calculations, the model is limited to the three genes implicated in azole resistance in the literature: *ERG11, ERG3*, and *ERG6*. Various point mutations in the *ERG11* gene were shown to alter the binding ability of azoles to the C14α-demethylase enzyme [8], [9]. Deletion of *ERG3* gene resulted in the accumulation of the toxic sterol precursor 14α-methylfecosterol, instead of the toxic sterol, allowing *C. albicans* survival [7], [10]. Disruption in the *ERG6* gene was shown to cause hypersusceptible strains to antifungal agents, but not to azoles [11]. Moreover, Akins (2005) noted that the *ERG6* gene is an attractive target for antifungals [12]. Hence, we deem it worth investigating whether it has an additive effect with fluconazole or not.

Another limitation of the study is that our simulations are limited to testing drug targets as an adjunct to fluconazole, since the model calibration was based on the study by Kelly et al. (1997), which compares the sterol compositions of the *C. albicans*, before and after fluconazole treatment.

The model uses ordinary differential equations (ODE) with mass-action kinetics, as first described in [13], which has since then been a standard method for mathematical modeling. The ODEs for the model without azole (normal pathway) are shown in Equation 1, as:

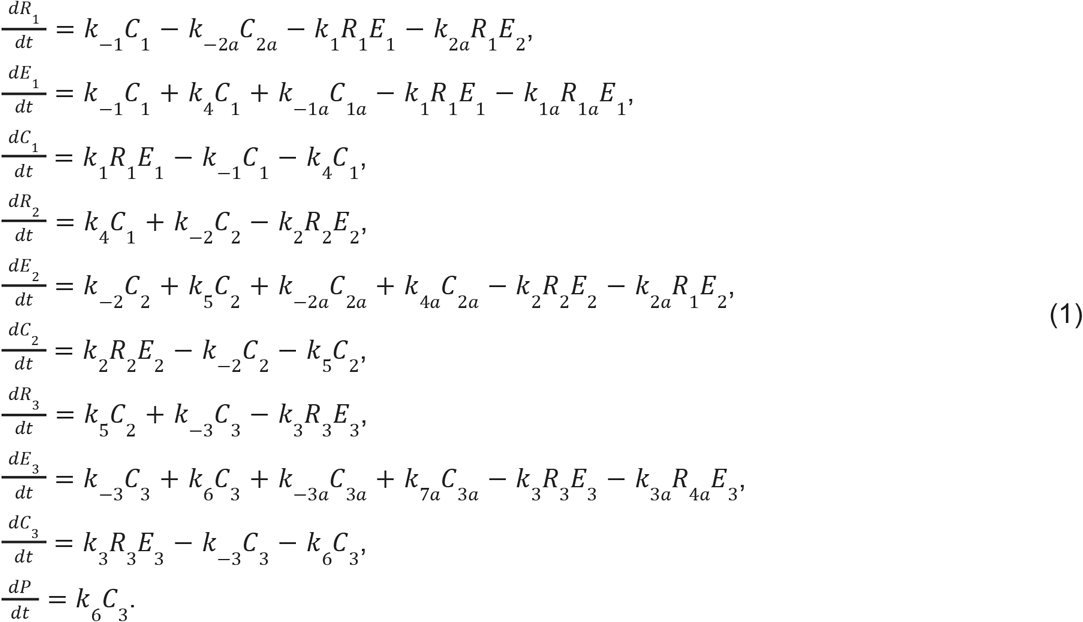

Whereas, the ODEs for the model with azole (alternative pathway) are shown in Equation 2, as:

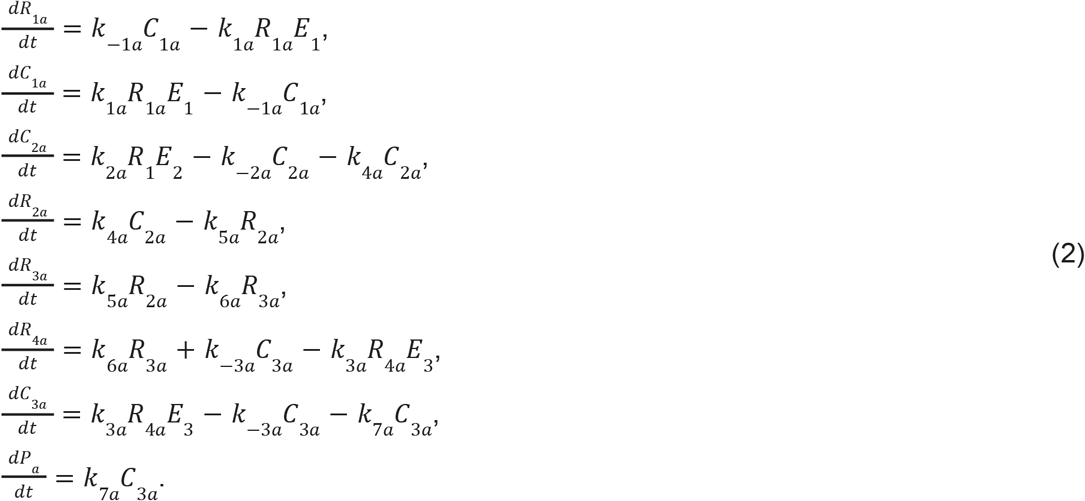

The constraint equations are shown in Equation 3, as:

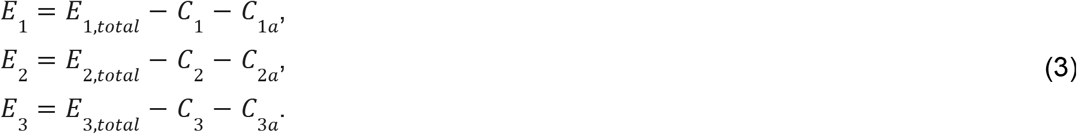

In the study by Hargrove et al. (2017), they used a molar ratio of 1:2:50 for enzyme, inhibitor, and substrate, respectively [14]. We tested it in our model and observed that the enzyme concentration was insufficient for the inhibitor to fully take its effect. As such, we tweaked the assignment of the molar ratio to 4:2:50 for enzyme, inhibitor, and substrate, respectively. The fluconazole concentration is set at 16 μg/ml, following Kelly et al. (1997). The parameter values, shown in Table 1, were obtained by calibrating the model to the sterol compositions described by Kelly et al. (1997), such that in a fluconazole-sensitive *C. albicans* strain, it is mainly composed of ergosterol (98%), and the rest is classified as unknown (2%). We distribute the unknown 2% to the remaining metabolites in the pathway. Whereas, in a fluconazole-resistant strain, the sterol composition is as follows: ergosterol (2%), eburicol (16.1%), obtusifoliol (34.5%), 14α-methyl-3,6-diol (45.2%), and unknown (2.2%). We again distribute the unknown 2.2% to the remaining metabolites in the pathway (see Table 2 in [10]). Following Kelly et al. (1997), the simulations are run for 24 hours, and the pertinent results are tabulated thereafter.

**Table 1.**
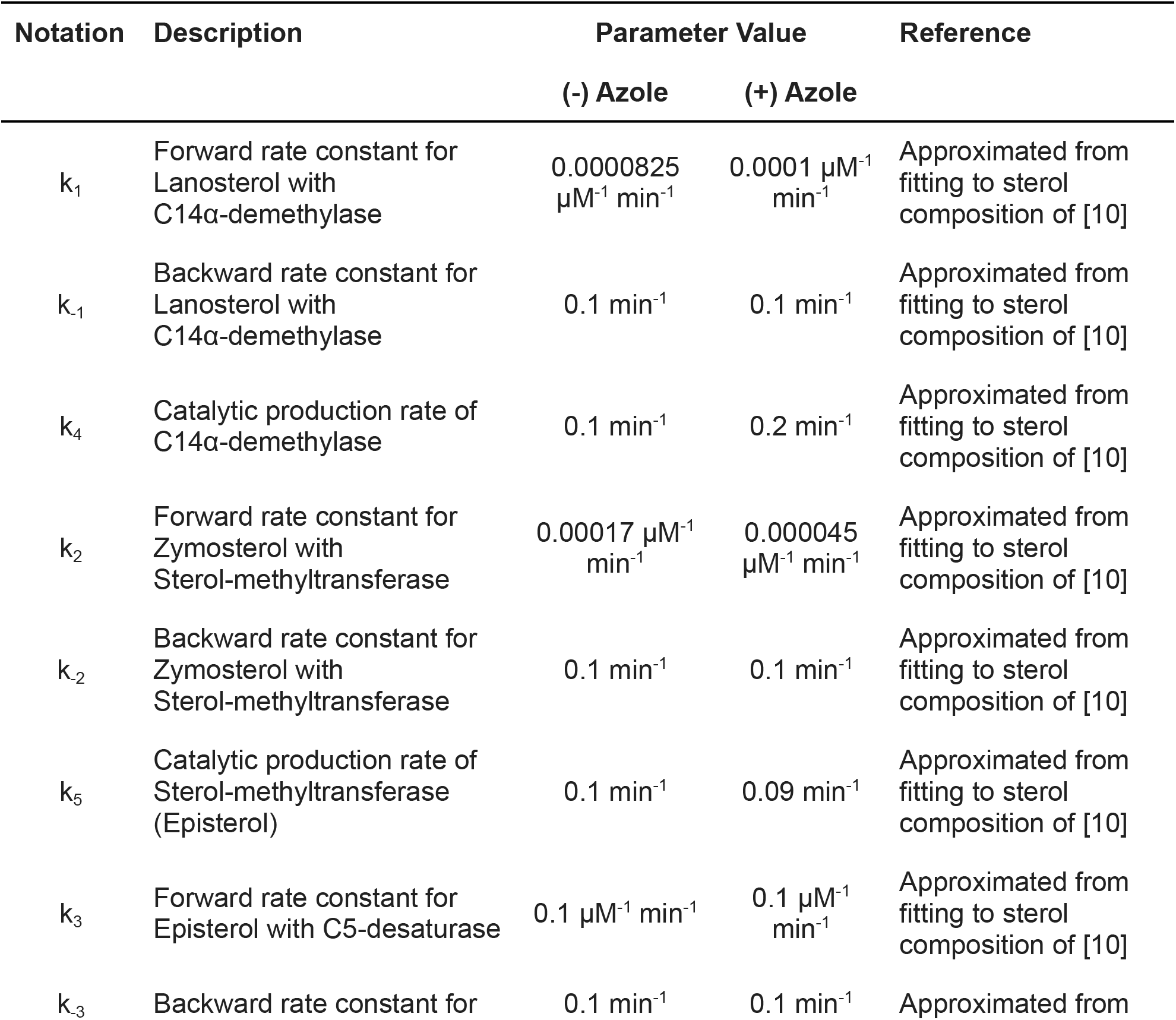
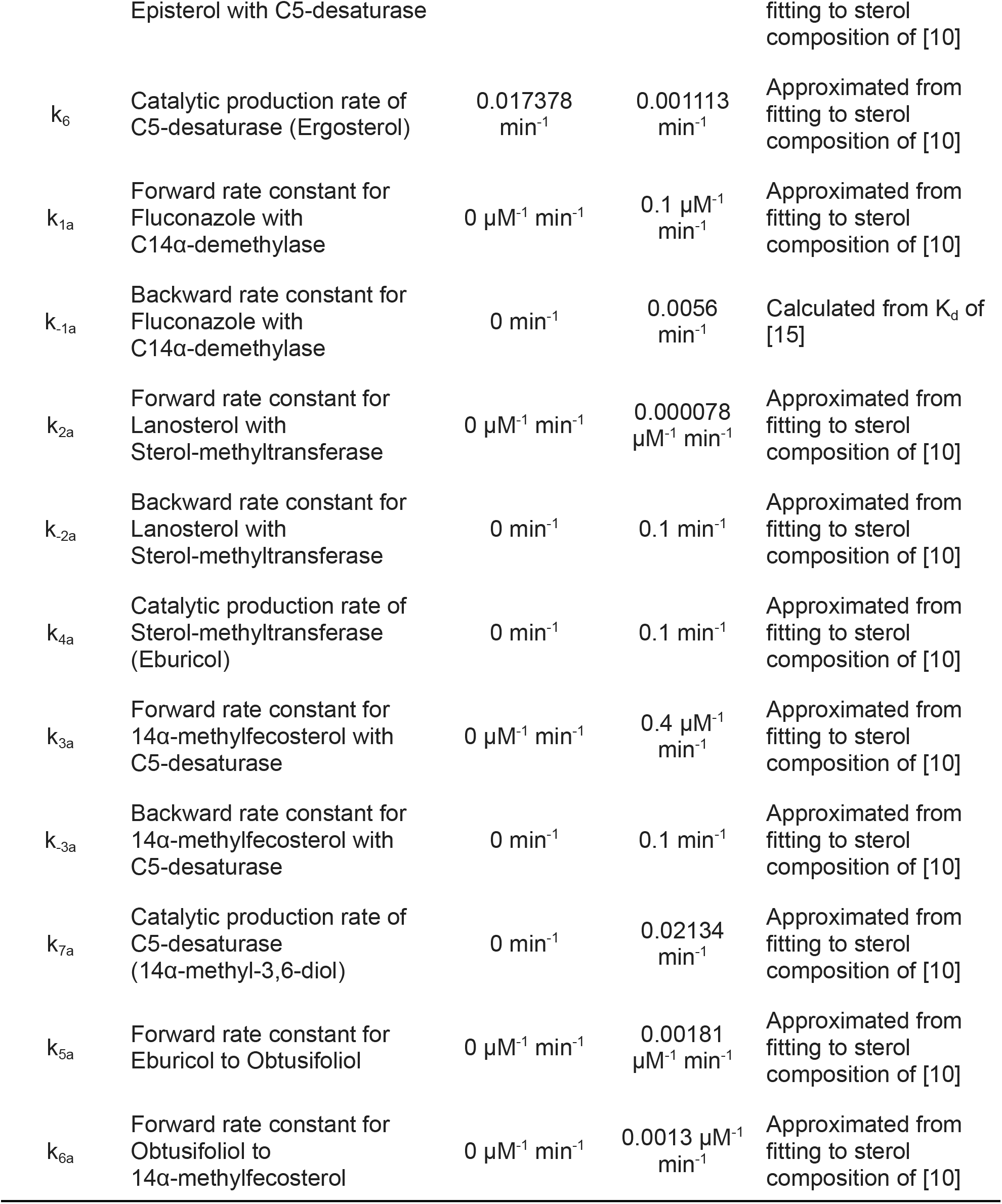
Model parameters

**Table 2.**
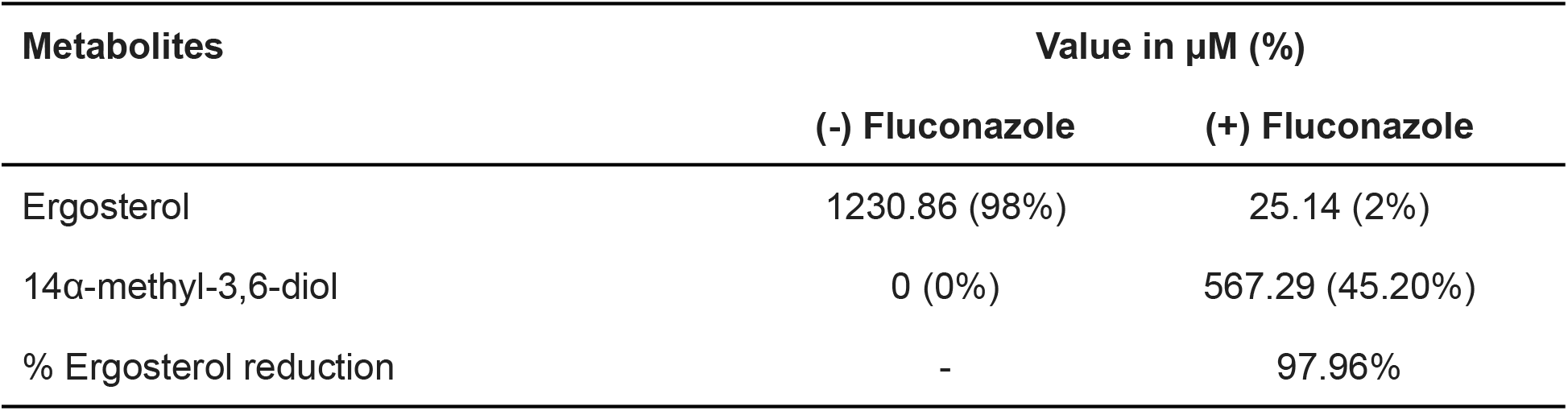
Pertinent results of the model calibration to Kelly et al. (1997)

| Metabolites | Value in $\mu\text{M}$ (%) | |
| --- | --- | --- |
|  | (-) Fluconazole | (+) Fluconazole |
| Ergosterol | 1230.86 (98%) | 25.14 (2%) |
| 14α-methyl-3,6-diol | 0 (0%) | 567.29 (45.20%) |
| % Ergosterol reduction | - | 97.96% |

The model generated the plots shown in Figure S1. The pertinent results are shown in Table 2.

The clinical breakpoints as determined from the 24-hour Clinical & Laboratory Standards Institute (CLSI) broth microdilution method for fluconazole against *C. albicans* are as follows: Susceptible (minimum inhibitory concentration (MIC) ≤ 2 μg/ml), Dose-dependent (MIC = 4 μg/ml), and Resistant (MIC ≥ 8 μg/ml) [16]. We re-calibrated our model to accommodate the MIC breakpoint for fluconazole susceptibility (MIC = 2 μg/ml) to determine its threshold for fluconazole susceptibility. To compute for the Molar concentration of 2 μg/ml, we use the following calculation: (molecular weight of fluconazole: 306.3g/mol from [17])

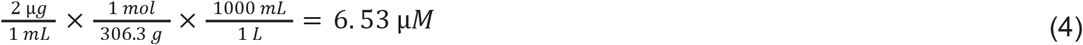

The re-calibrated model generated the plots shown in Figure S2. This will serve as the baseline for the simulations in Section 3. The pertinent results are shown in Table 3.

**Table 3.** Pertinent results of the model re-calibration to the MIC breakpoint for fluconazole susceptibility

| Metabolites | Value in $\mu\text{M}$ (%) | |
| --- | --- | --- |
|  | (-) Fluconazole | (+) Fluconazole |
| Ergosterol | 1230.95 (98%) | 65.41 (5.22%) |
| 14α-methyl-3,6-diol | 0 (0%) | 222.01 (17.73%) |
| % Ergosterol reduction | - | 94.69% |

In experimental studies, antifungal susceptibility is quantified by visual spotting assay on YEPD medium, such as in [5], disc diffusion method, such as in [8], or broth microdilution method, such as in [18]. However, these methods are not feasible to use in mathematical models, since they cannot be quantified. As such, we look for quantitative methods at the metabolite level to circumvent this limitation.

Arthington-Skaggs et al. (1999) proposes the use of percent ergosterol reduction as a quantitative method to evaluate antifungal susceptibility [19]. However, Kelly et al. (1997) found that an accumulation of the toxic sterol causes the growth arrest of the *C. albicans* [10]. Another interesting result found by Martel et al. (2010) is that antifungal susceptibility holds when the dominant fraction of sterol is ergosterol, regardless of the amount of the toxic sterol [7].

Considering the above methods for antifungal susceptibility, we propose the following criteria based on the data gathered in Table 3 to determine the threshold for fluconazole resistance in *C. albicans*:

1. when the % toxic sterol (14α-methyl-3,6-diol) is less than 17.73%;
2. when the % ergosterol reduction is less than 94.69%;
3. when ergosterol is the dominant fraction among the metabolites.

When all 3 criteria are satisfied, we remark it as good fluconazole resistance. When 2 out of 3 are satisfied, we denote it as fair fluconazole resistance. When only 1 out of 3 is satisfied, we deem it as an equivocal result, such that no conclusion can be drawn. Conversely, when all criteria are not satisfied, we remark it as a good target for antifungals as adjunct to fluconazole.

## 3. Results

The model was run to simulate several scenarios to investigate possible drug targets in the ergosterol pathway in *C. albicans*.

### 3.1 Variations in lanosterol concentration

The lanosterol concentration was varied as follows: 0x, 0.01x, 0.1x, 1x, 10x, 100x the baseline value (see Figure S3), represented by case 1 to 6, respectively. Case 1 (0x) simulates a total inhibition of lanosterol. It results in the demise of the *C. albicans*, since no ergosterol was produced, which is essential for yeast survival. Cases 2 (0.01x) and 3 (0.1x) simulate the partial inhibition of lanosterol. They yield a higher percentage of the toxic sterol, and ergosterol as the dominant metabolite, but a lower percentage of ergosterol reduction, satisfying 2 out of 3 criteria, which denotes a fair fluconazole resistance. Case 4 (1x) denotes the default case, which serves as our baseline, as shown in Table 3. Cases 5 (10x) and 6 (100x) simulate the overproduction of lanosterol. They yield a lower percentage of the toxic sterol, but a higher percentage of ergosterol reduction, satisfying 1 out of 3 criteria, which denotes an equivocal result. Table 4 summarizes the results pertinent to the criteria for fluconazole resistance.

**Table 4.**
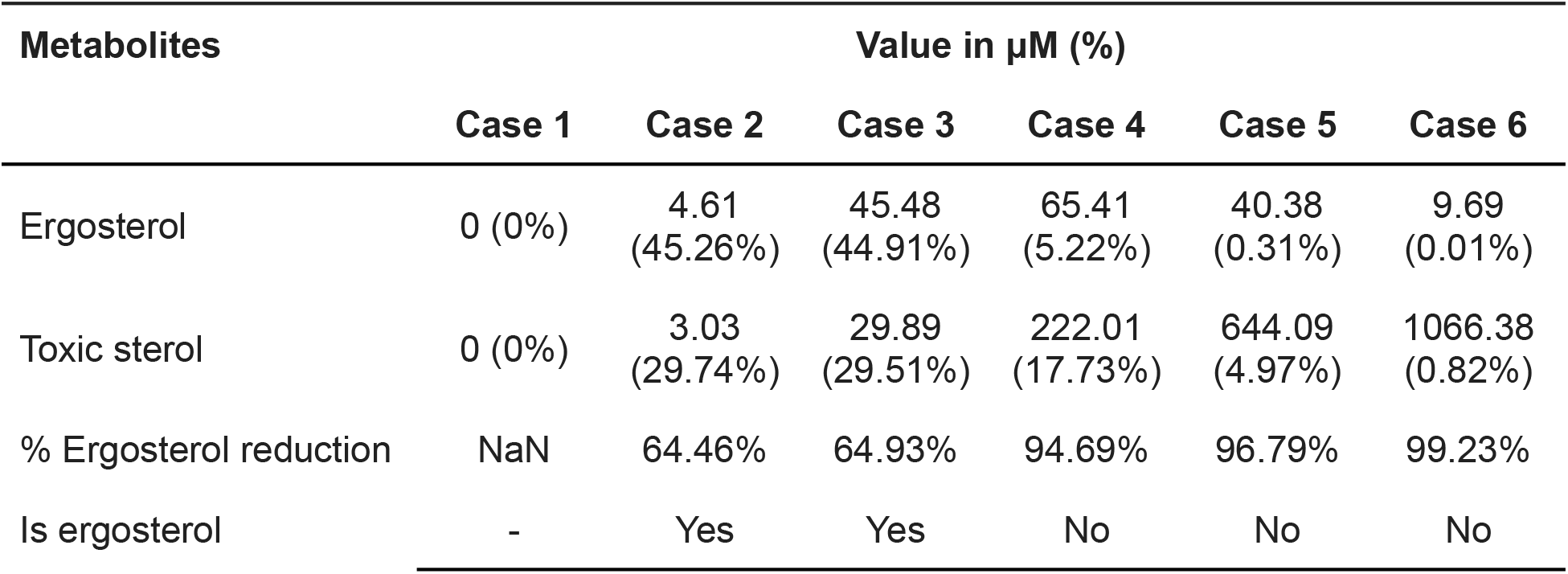
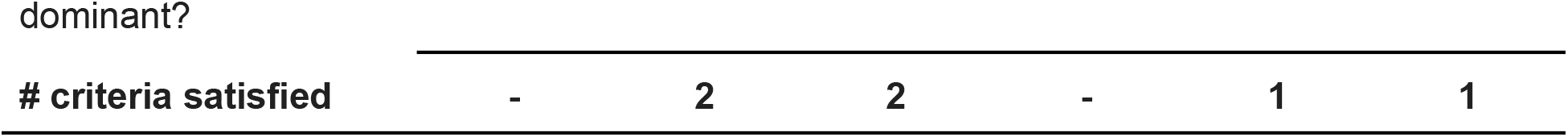
Pertinent results of variations in lanosterol concentration simulation

| Metabolites | Value in $\mu\text{M}$ (%) | | | | | |
| --- | --- | --- | --- | --- | --- | --- |
|  | Case 1 | Case 2 | Case 3 | Case 4 | Case 5 | Case 6 |
| Ergosterol | 0 (0%) | 4.61<br>(45.26%) | 45.48<br>(44.91%) | 65.41<br>(5.22%) | 40.38<br>(0.31%) | 9.69<br>(0.01%) |
| Toxic sterol | 0 (0%) | 3.03<br>(29.74%) | 29.89<br>(29.51%) | 222.01<br>(17.73%) | 644.09<br>(4.97%) | 1066.38<br>(0.82%) |
| % Ergosterol reduction | NaN | 64.46% | 64.93% | 94.69% | 96.79% | 99.23% |
| Is ergosterol | - | Yes | Yes | No | No | No |
| **# criteria satisfied** | - | 2 | 2 | - | 1 | 1 |

### 3.2 Variations in C14α-demethylase concentration (Erg11p)

The C14α-demethylase enzyme concentration was varied as follows: 0x, 0.01x, 0.1x, 1x, 10x, 100x the baseline value (see Figure S4), represented by case 1 to 6, respectively. Case 1 (0x) simulates a total inhibition of C14α-demethylase. This results in the demise of the *C. albicans*, since no ergosterol was produced, which is essential for yeast survival. Cases 2 (0.01x) and 3 (0.1x) simulate the partial inhibition of C14α-demethylase. They yield a higher percentage of the toxic sterol, and a higher percentage of ergosterol reduction, satisfying 0 out of 3 criteria, which denotes a good target for antifungals as adjunct to fluconazole. Case 4 (1x) denotes the default case, which serves as our baseline, as shown in Table 3. Cases 5 (10x) and 6 (100x) simulate the overexpression of the *ERG11* gene. They yield a lower percentage of the toxic sterol, and a lower percentage of ergosterol reduction, satisfying 2 out of 3 criteria, which denotes a fair fluconazole resistance. Table 5 summarizes the results pertinent to the criteria for fluconazole resistance.

**Table 5.** Pertinent results of variations in C14α-demethylase concentration simulation

| Metabolites | Value in μM (%) | | | | | |
| --- | --- | --- | --- | --- | --- | --- |
|  | Case 1 | Case 2 | Case 3 | Case 4 | Case 5 | Case 6 |
| Ergosterol | 0 (0%) | 0.11 (0.01%) | 3.43 (0.27%) | 65.41 (5.22%) | 79.60 (6.36%) | 81.95 (6.55%) |
| Toxic sterol | 589.71 (46.17%) | 589.63 (46.17%) | 587.18 (46.08%) | 222.01 (17.73%) | 31.16 (2.49%) | 3.64 (0.29%) |
| % Ergosterol reduction | NaN | 99.68% | 98.91% | 94.69% | 93.74% | 93.59% |
| Is ergosterol dominant? | - | No | No | No | No | No |
| **# criteria satisfied** | - | 0 | 0 | - | 2 | 2 |

### 3.3 Variations in fluconazole concentration

The fluconazole concentration was varied as follows: 0x, 0.01x, 0.1x, 1x, 10x, 100x the baseline value (see Figure S5), represented by case 1 to 6, respectively. Case 1 (0x) simulates a situation where no fluconazole was given. This results in the proliferation of the *C. albicans*, since the dominant metabolite is ergosterol, which is essential for yeast survival. Cases 2 (0.01x) and 3 (0.1x) simulate giving a fluconazole with a dose lower than its minimum inhibitory concentration (MIC), i.e. 2 μg/ml. They yield a lower percentage of the toxic sterol, and a lower percentage of ergosterol reduction, satisfying 2 out of 3 criteria, which denotes a fair fluconazole resistance. Case 4 (1x) denotes the default case, which serves as our baseline, as shown in Table 3. Cases 5 (10x) and 6 (100x) simulate giving fluconazole with a dose higher than its MIC. They yield a higher percentage of the toxic sterol, and a higher percentage of ergosterol reduction, satisfying 0 out of 3 criteria, which denotes a good target for antifungals as adjunct to fluconazole. Table 6 summarizes the results pertinent to the criteria for fluconazole resistance.

**Table 6.** Pertinent results of variations in fluconazole concentration simulation

| Metabolites | Value in $\mu\text{M}$ (%) | | | | | |
| --- | --- | --- | --- | --- | --- | --- |
|  | Case 1 | Case 2 | Case 3 | Case 4 | Case 5 | Case 6 |
| Ergosterol | 1230.86<br>(98%) | 66.73<br>(5.33%) | 66.62<br>(5.32%) | 65.41<br>(5.22%) | 3.89<br>(0.31%) | 0.11<br>(0.01%) |
| 14a-methyl-3,6-diol | 0 (0%) | 204.17<br>(16.31%) | 205.67<br>(16.43%) | 222.01<br>(17.73%) | 586.84<br>(46.06%) | 589.62<br>(46.17%) |
| % Ergosterol reduction | 0% | 94.58% | 94.59% | 94.69% | 99.68% | 99.99% |
| Is ergosterol dominant? | - | No | No | No | No | No |
| # criteria satisfied | - | 2 | 2 | - | 0 | 0 |

### 3.4 Variation in sterol-methyltransferase concentration (Erg6p)

The sterol-methyltransferase enzyme concentration was varied as follows: 0x, 0.01x, 0.1x, 1x, 10x, 100x the baseline value (see Figure S6), represented by case 1 to 6, respectively. Case 1 (0x) simulates a total inhibition of sterol-methyltransferase. This results in the demise of the *C. albicans*, since no ergosterol was produced, which is essential for yeast survival. Cases 2 (0.01x) and 3 (0.1x) simulate the partial inhibition of sterol-methyltransferase. They yield a lower percentage of the toxic sterol, and a lower percentage of ergosterol reduction, satisfying 2 out of 3 criteria, which denotes a fair fluconazole resistance. Case 4 (1x) denotes the default case, which serves as our baseline, as shown in Table 3. Cases 5 (10x) and 6 (100x) simulate the overexpression of the *ERG6* gene. They yield a higher percentage of the toxic sterol, and a higher percentage of ergosterol reduction, satisfying 0 out of 3 criteria, which denotes a good target for antifungals as adjunct to fluconazole. Table 7 summarizes the results pertinent to the criteria for fluconazole resistance.

**Table 7.** Pertinent results of variations in sterol-methyltransferase concentration simulation

| Metabolites | Value in $\mu\text{M}$ (%) | | | | | |
| --- | --- | --- | --- | --- | --- | --- |
|  | Case 1 | Case 2 | Case 3 | Case 4 | Case 5 | Case 6 |
| Ergosterol | 0 (0%) | 9.31<br>(0.72%) | 58.54<br>(4.67%) | 65.41<br>(5.22%) | 46.52<br>(3.71%) | 15.07<br>(1.18%) |
| Toxic sterol | 0 (0%) | 3.63<br>(0.28%) | 34.58<br>(2.76%) | 222.01<br>(17.73%) | 660.83<br>(52.70%) | 796.76<br>(62.32%) |
| % Ergosterol reduction | NaN | 77.83% | 85.13% | 94.69% | 96.32% | 98.81% |
| Is ergosterol dominant? | - | No | No | No | No | No |
| **# criteria satisfied** | - | **2** | **2** | - | **0** | **0** |

### 3.5 Variation in C5-desaturase concentration (Erg3p)

The C5-desaturase enzyme concentration was varied as follows: 0x, 0.01x, 0.1x, 1x, 10x, 100x the baseline value (see Figure S7), represented by case 1 to 6, respectively. Case 1 (0x) simulates a total inhibition of C5-desaturase. This results in the demise of the *C. albicans*, since no ergosterol was produced, which is essential for yeast survival. Cases 2 (0.01x) and 3 (0.1x) simulate the partial inhibition of C5-desaturase. They yield a lower percentage of the toxic sterol, but a higher percentage of ergosterol reduction, satisfying 1 out of 3 criteria, which denotes an equivocal result. Case 4 (1x) denotes the default case, which serves as our baseline, as shown in Table 3. Cases 5 (10x) and 6 (100x) simulate the overexpression of the *ERG3* gene. They yield a higher percentage of the toxic sterol, but a lower percentage of ergosterol reduction and ergosterol as the dominant metabolite, satisfying 2 out of 3 criteria, which denotes a fair fluconazole resistance. Table 8 summarizes the results pertinent to the criteria for fluconazole resistance.

**Table 8.** Pertinent results of variations in C5-desaturase concentration simulation

| Metabolites | Value in $\mu\text{M}$ (%) | | | | | |
| --- | --- | --- | --- | --- | --- | --- |
|  | Case 1 | Case 2 | Case 3 | Case 4 | Case 5 | Case 6 |
| Ergosterol | 0 (0%) | 0.51 (0.04%) | 5.37 (0.41%) | 65.41 (5.22%) | 431.35 (44.26%) | 432.11 (44.34%) |
| Toxic sterol | 0 (0%) | 5.87 (0.45%) | 52.20 (4.02%) | 222.01 (17.73%) | 266.99 (27.39%) | 267.00 (27.40%) |
| % Ergosterol reduction | NaN | 96.03% | 95.81% | 94.69% | 66.36% | 66.31% |
| Is ergosterol dominant? | - | No | No | No | Yes | Yes |
| **# criteria satisfied** | - | **1** | **1** | - | **2** | **2** |

## 4. Discussion

The simulation for variations in lanosterol concentration showed that its total inhibition, shown in Table 4 case 1, by *ERG1* gene deletion results in the demise of the *C. albicans* due to its lack of ergosterol. This is in agreement with the study by Pasrija et al. (2005) that *erg1* mutant showed a total lack of ergosterol, hence leading to increased azole susceptibility [20]. However, its partial inhibition, shown in Table 4 cases 2 & 3, results in a fair fluconazole resistance. An example of this are allylamine antifungals, which inhibit the squalene epoxidase enzyme in the pathway upstream of lanosterol. However, Kane and Carter showed that there is a synergistic effect when combining terbinafine (under allylamines) and fluconazole against *C. albicans* [21]. Allylamines cause fungal cell death by depleting the ergosterol content, and by accumulation of squalene [22]. The discrepancy may be due to our model being limited to the three implicated genes: *ERG11, ERG3*, and *ERG6* genes. Moreover, an overproduction of lanosterol, as shown in Table 4 cases 5 & 6, yields an equivocal result, hence no conclusion can be drawn. This indicates the need for further experiments, or further extension of the model to include upstream pathways, such as the squalene epoxidase enzyme.

The simulation for variations in C14α-demethylase concentration confirmed that its partial inhibition, as shown in Table 5 cases 2 & 3, is a good target of antifungals, as it should be, since it has been known that this enzyme is the target of azole drugs, such as fluconazole, leading to the demise of the *C. albicans* [23], [24]. On the other hand, an overexpression of the *ERG11* gene is a known azole resistance mechanism [25]. This has been confirmed by our simulation, shown in Table 5 cases 5 and 6, such that there was a lower percentage of the toxic sterol, and a lower percentage of ergosterol reduction.

The simulation for variations in fluconazole concentration confirmed that the effect of fluconazole in *C. albicans* is dose-dependent, such that doses lower than its minimum inhibitory concentration (MIC) do not fully eradicate the fungi, whereas doses greater than its MIC eradicates them. Hence, it is crucial to have an accurate quantitative method to determine the MIC for antifungal susceptibility testing [16].

The simulation for variations in sterol-methyltransferase concentration showed that its partial inhibition, as shown in Table 7 cases 2 & 3, yields a fair fluconazole resistance. This agrees with the findings of Kelly et al. (1997) that a defect in the *ERG6* gene leads to toxic sterol accumulation, hence the resistance to fluconazole [10]. This resistance mechanism is also described in the review by Bhattacharya et al. (2020) [24]. On the other hand, our results showed that an overexpression of the *ERG6* gene, shown in Table 7 cases 5 & 6, is a good target for antifungals as adjunct to fluconazole. Bhattacharya et al. (2018) noted that in *Saccharomyces cerevisiae*, an overexpression of the *ERG6* gene encourages the initiation of the alternative pathway, leading to the production of toxic sterol [26]. In the review by Bhattacharya et al. (2020), the *ERG6* gene contributes to the formation of the toxic sterol, in the presence of azoles [24]. Hence, we can infer that an overexpression of the *ERG6* gene can lead to an increase in the formation of the toxic sterol, resulting in hypersusceptibility to azoles. As such, this can have an additive effect when combined with fluconazole to circumvent the resistance by *C. albicans* against it. The overexpression of the *ERG6* gene yields a higher percentage of the toxic sterol, and a higher percentage of ergosterol reduction. Hence, we recommend doing further experiments on the sterol-methyltransferase enzyme to confirm that its overexpression is a possible drug target as an adjunct to fluconazole, to validate our simulation result.

The simulation for variations in C5-desaturase concentration showed that its partial inhibition, as shown in Table 8 cases 2 & 3, leads to an equivocal result, hence no conclusion can be drawn. Although in the literature, it is known that its mutation is linked with azole resistance [5], [7], [24]. On the other hand, our simulation of an overexpression of the *ERG3* gene, shown in Table 8 cases 5 & 6, results in a fair fluconazole resistance. No studies have been done yet regarding this in the literature, however our results showed that it yields a lower percentage of ergosterol reduction and ergosterol as the dominant metabolite. Hence, we recommend against looking into this enzyme as a possible drug target as an adjunct to fluconazole.

## 5. Conclusions

From the model simulations, we found the following results: (1) a partial inhibition of sterol-methyltransferase yields a fair amount of fluconazole resistance, (2) an overexpression of the *ERG6* gene, leading to increased sterol-methyltransferase enzyme, is a good target of antifungals as an adjunct to fluconazole, (3) a partial inhibition of lanosterol yields a fair amount of fluconazole resistance, (4) C5-desaturase is not a good target of antifungals as an adjunct to fluconazole, (5) C14α-demethylase is confirmed to be a good target for fluconazole, and (6) the dose-dependent effect of fluconazole is confirmed. This study hopes to aid experimenters narrow down the possible drug targets prior to doing costly and time-consuming experiments, and to serve as a cross-validation tool for experimental data.

We recommend extending the model to include other genes involved in the ergosterol biosynthesis pathway, such as the *ERG1* gene that expresses squalene epoxidase, which is the target of allylamine antifungals, or to include other mechanisms involved in azole resistance mechanism, such as the *CDR1, CDR2*, and *MDR1* genes, which are involved in the drug efflux transporter. We also recommend doing further experiments on the sterol-methyltransferase enzyme to confirm that its overexpression is a possible target for antifungals as an adjunct to fluconazole, to validate our simulation result.

## Supplementary Materials

Figure S1: Plots generated by the model calibrated from Kelly et al., (1997); Figure S2: Plots generated by the model re-calibrated to the minimum inhibitory concentration (MIC) breakpoint for fluconazole susceptibility; Figure S3: Plots generated by the simulation of variations in lanosterol concentration; Figure S4: Plots generated by the simulation of variations in C14α-demethylase concentration; Figure S5. Plots generated by the simulation of variations in fluconazole concentration; Figure S6: Plots generated by the simulation of variations in sterol-methyltransferase concentration; Figure S7: Plots generated by the simulation of variations in C5-desaturase concentration.

## Author Contributions

Conceptualization, P.K.Y., L.S.M, A.R.L.; methodology, P.K.Y, L.S.M, A.R.L.; software, P.K.Y.; validation, P.K.Y., L.S.M., A.R.L.; formal analysis, P.K.Y., L.S.M., A.R.L.; investigation, P.K.Y., L.S.M., A.R.L.; resources, P.K.Y., L.S.M., A.R.L.; data curation, P.K.Y., L.S.M., A.R.L.; writing—original draft preparation, P.K.Y.; writing—review and editing, P.K.Y., L.S.M., A.R.L.; visualization, P.K.Y.; supervision, L.S.M., A.R.L.; project administration, P.K.Y., L.S.M., A.R.L.; funding acquisition, P.K.Y., L.S.M., A.R.L. All authors have read and agreed to the published version of the manuscript.

## Institutional Review Board Statement

Not applicable.

## Informed Consent Statement

Not applicable.

## Data Availability Statement

Not applicable.

## Acknowledgments

P.K.Y. would like to thank the Department of Science and Technology (DOST)-Science Education Institute (SEI) Career Incentive Program for the support. The authors would also like to thank the DOST-Philippine Council for Health Research and Development (PCHRD).

## Conflicts of Interes

The authors declare no conflict of interest.

