## Supplementary material for "Mathematical Modeling of Fluconazole Resistance in the Ergosterol Pathway of *Candida albicans*": Figure S1 - S7

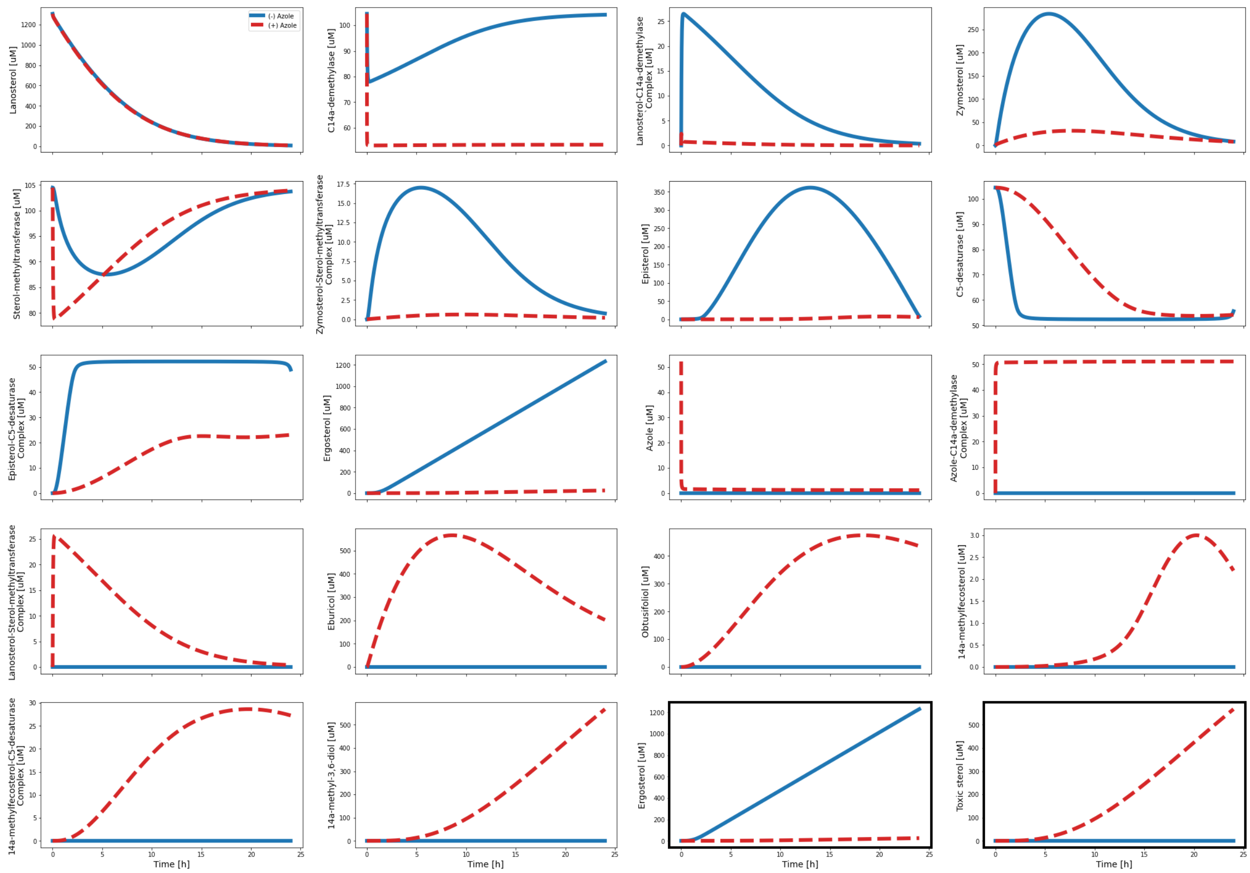


**Figure S1.** Plots generated by the model calibrated from Kelly et al., (1997). The normal pathway, when azole is absent, is denoted by blue solid lines. The alternative pathway, when azole is present, is denoted by red dashed lines. The x-axes denote the time in hours, while the y-axes represent the concentrations in μM. The last two subplots, emphasized in dark borders, reiterate the pertinent plots of ergosterol and the toxic sterol (14α-methyl-3,6-diol) for easier comparison.

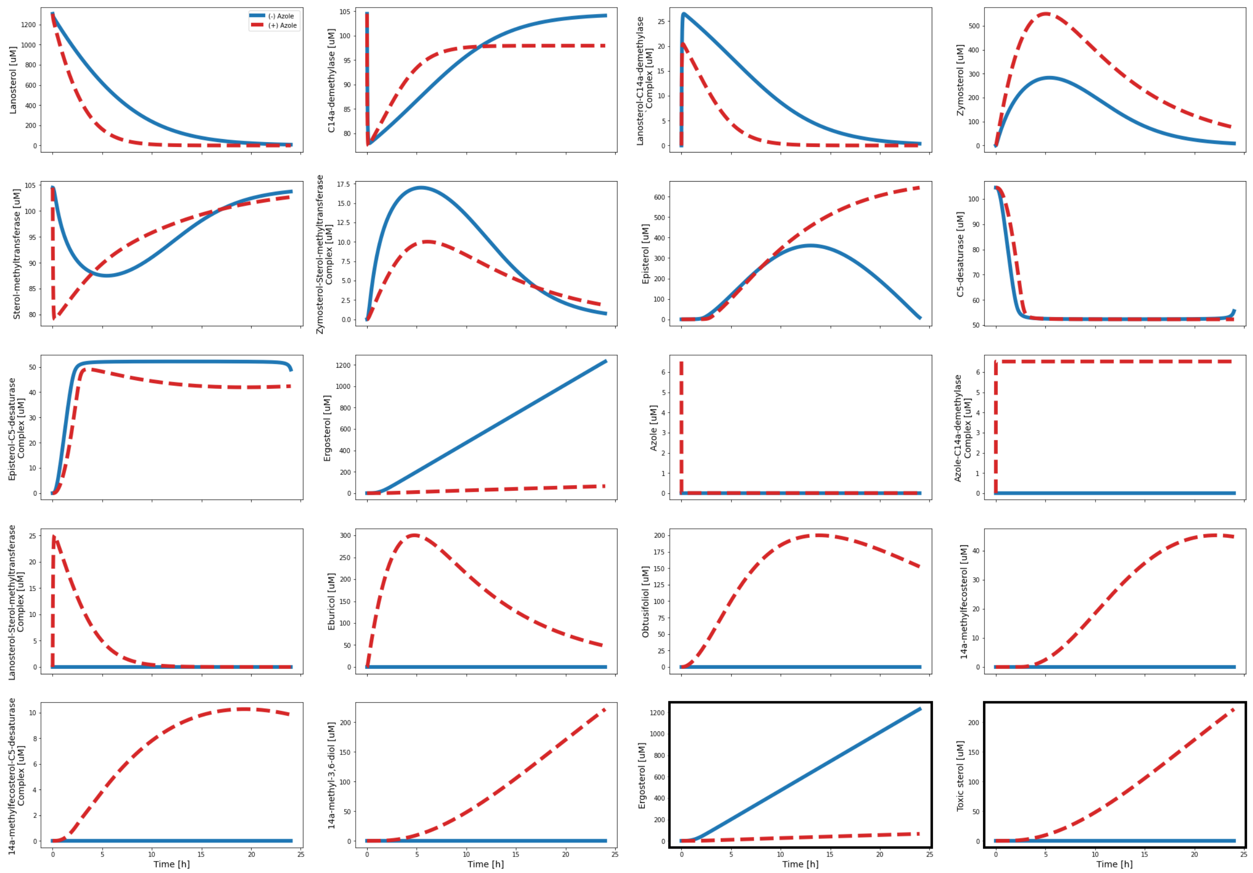


**Figure S2.** Plots generated by the model re-calibrated to the minimum inhibitory concentration (MIC) breakpoint for fluconazole susceptibility. The normal pathway, when azole is absent, is denoted by blue solid lines. The alternative pathway, when azole is present, is denoted by red dashed lines. The x-axes denote the time in hours, while the y-axes represent the concentrations in μM. The last two subplots, emphasized in dark borders, reiterate the pertinent plots of ergosterol and the toxic sterol (14α-methyl-3,6-diol) for easier comparison.

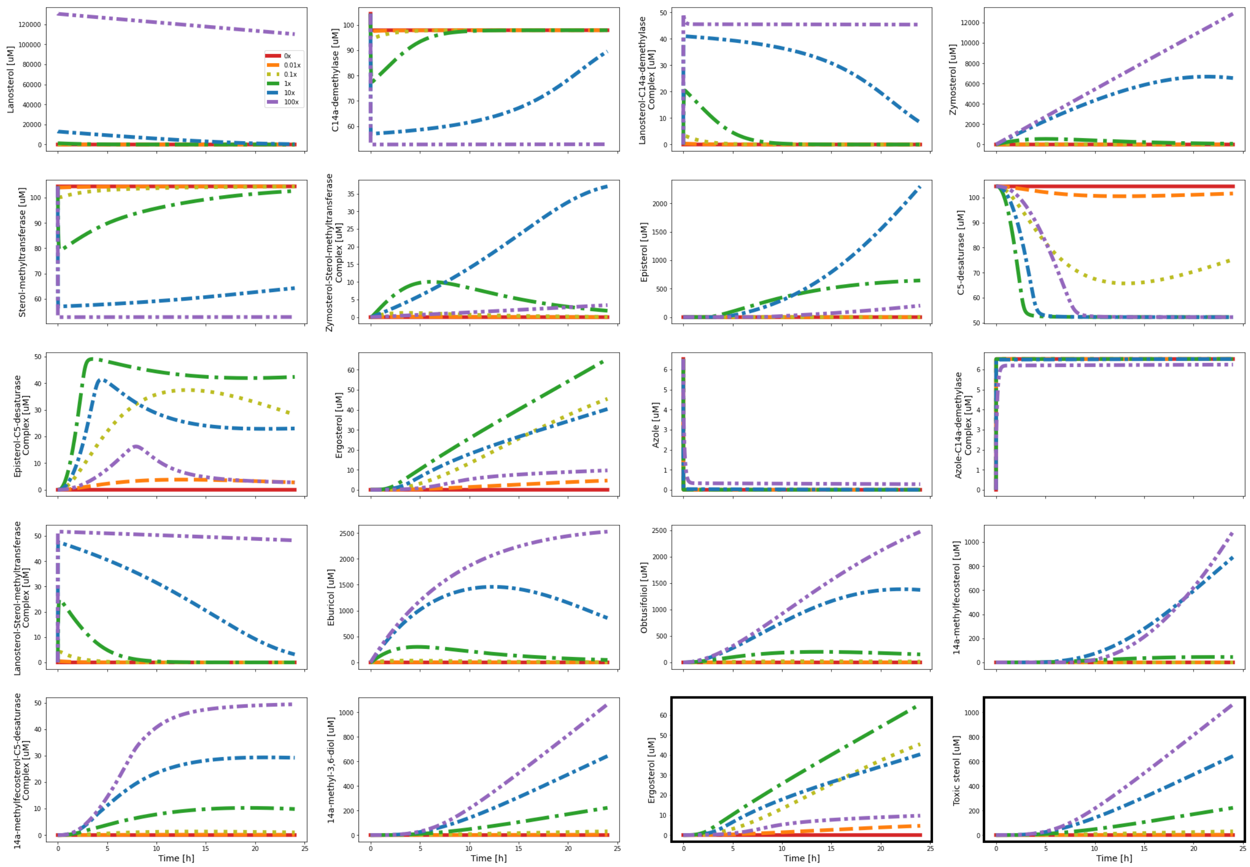


**Figure S3.** Plots generated by the simulation of variations in lanosterol concentration. The variations are as follows: 0x (red solid line), 0.01x (orange dashed line), 0.1x (yellow dotted line), 1x (green dashed dotted line), 10x (blue dense dashed dotted line), and 100x (purple dashed double dotted line). The x-axes denote the time in hours, while the y-axes represent the concentrations in µM. The last two subplots, emphasized in dark borders, reiterate the pertinent plots of ergosterol and the toxic sterol (14α-methyl-3,6-diol) for easier comparison.

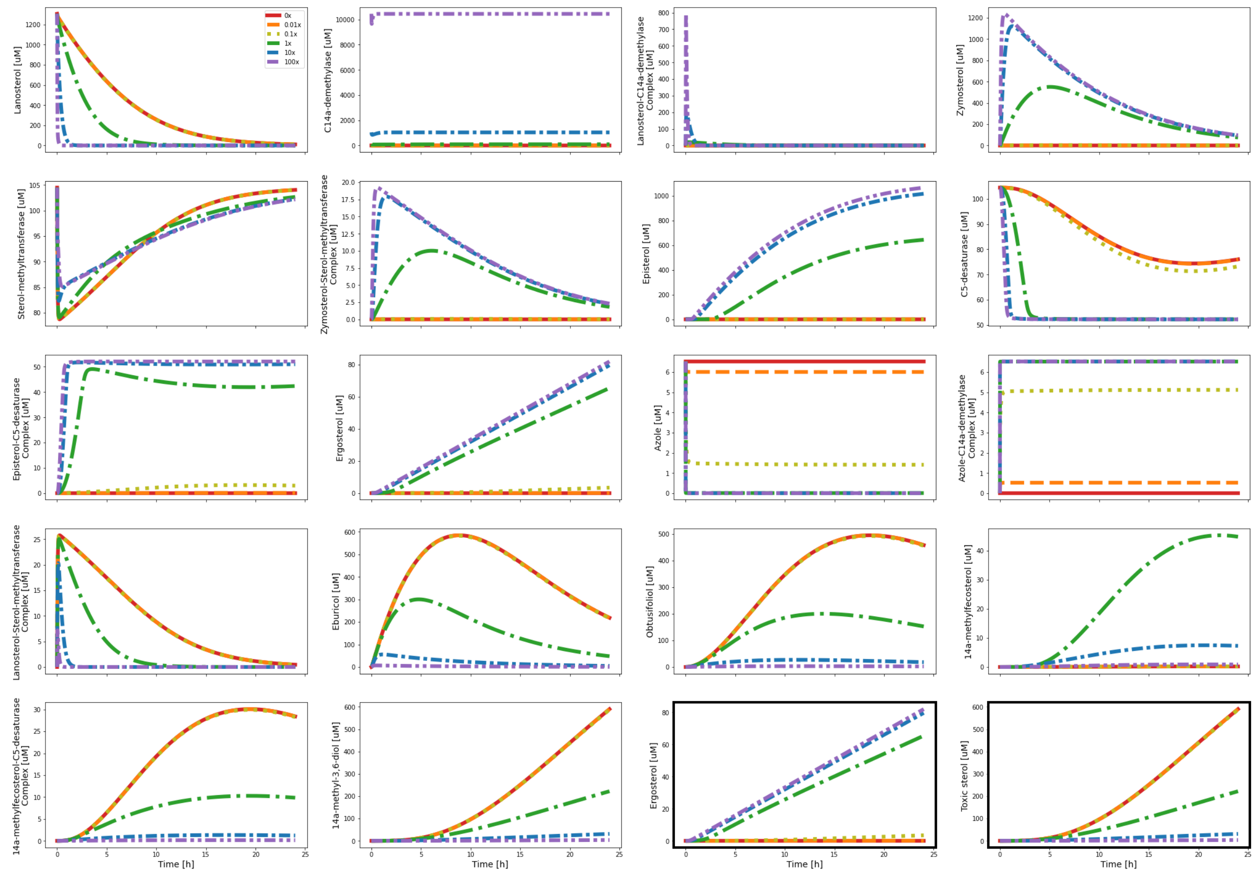


**Figure S4.** Plots generated by the simulation of variations in C14α-demethylase concentration. The variations are as follows: 0x (red solid line), 0.01x (orange dashed line), 0.1x (yellow dotted line), 1x (green dashed dotted line), 10x (blue dense dashed dotted line), and 100x (purple dashed double dotted line). The x-axes denote the time in hours, while the y-axes represent the concentrations in μM. The last two subplots, emphasized in dark borders, reiterate the pertinent plots of ergosterol and the toxic sterol (14α-methyl-3,6-diol) for easier comparison.

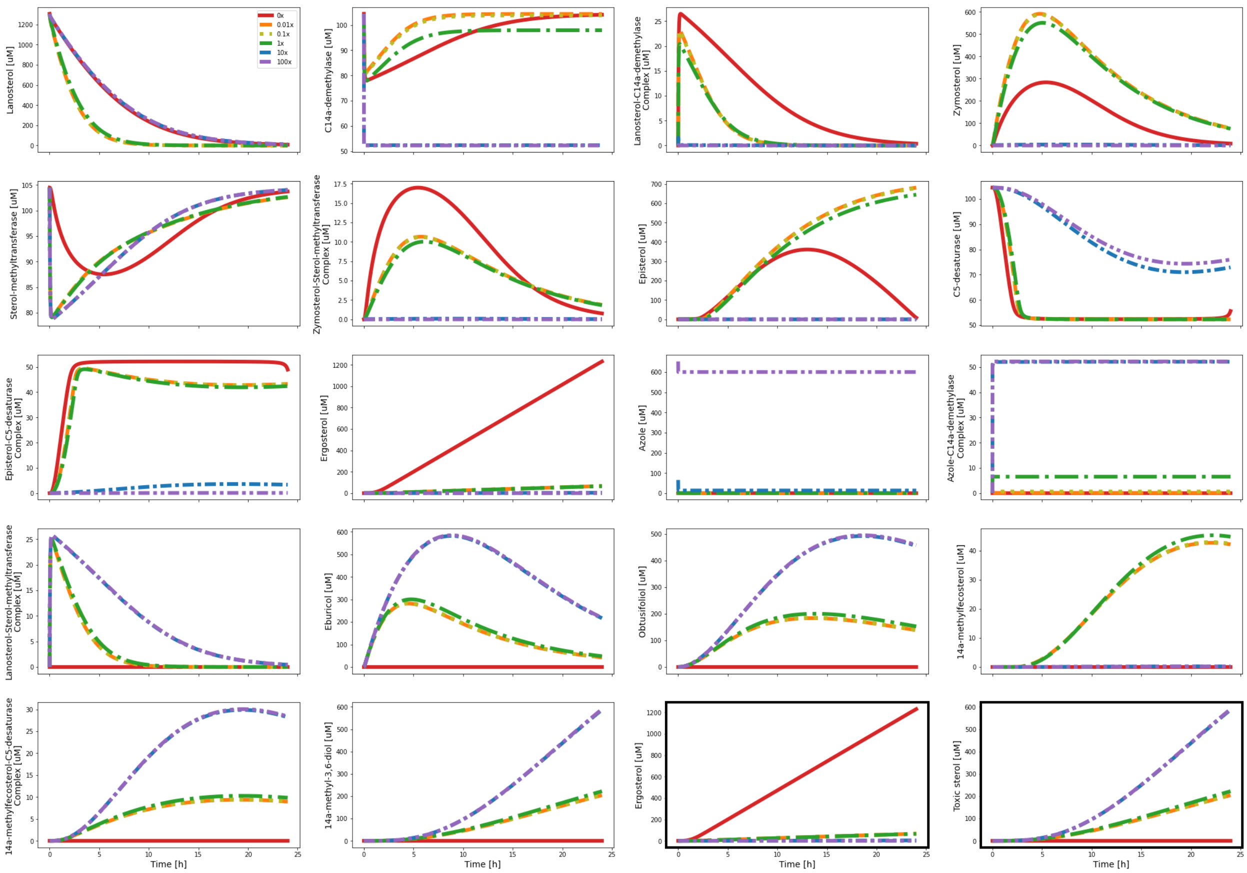


**Figure S5.** Plots generated by the simulation of variations in fluconazole concentration. The variations are as follows: 0x (red solid line), 0.01x (orange dashed line), 0.1x (yellow dotted line), 1x (green dashed dotted line), 10x (blue dense dashed dotted line), and 100x (purple dashed double dotted line). The x-axes denote the time in hours, while the y-axes represent the concentrations in μM. The last two subplots, emphasized in dark borders, reiterate the pertinent plots of ergosterol and the toxic sterol (14α-methyl-3,6-diol) for easier comparison.

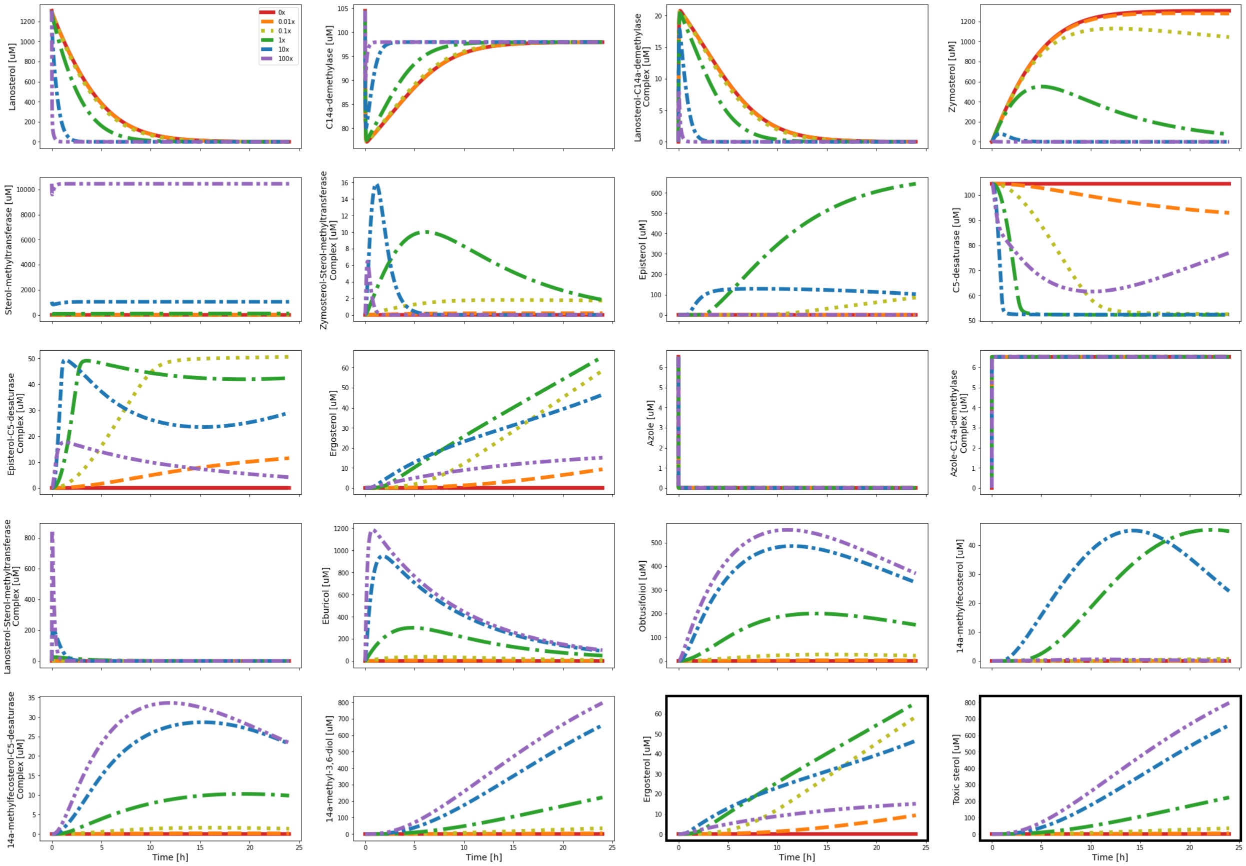


**Figure S6.** Plots generated by the simulation of variations in sterol-methyltransferase concentration. The variations are as follows: 0x (red solid line), 0.01x (orange dashed line), 0.1x (yellow dotted line), 1x (green dashed dotted line), 10x (blue dense dashed dotted line), and 100x (purple dashed double dotted line). The x-axes denote the time in hours, while the y-axes represent the concentrations in μM. The last two subplots, emphasized in dark borders, reiterate the pertinent plots of ergosterol and the toxic sterol (14α-methyl-3,6-diol) for easier comparison.

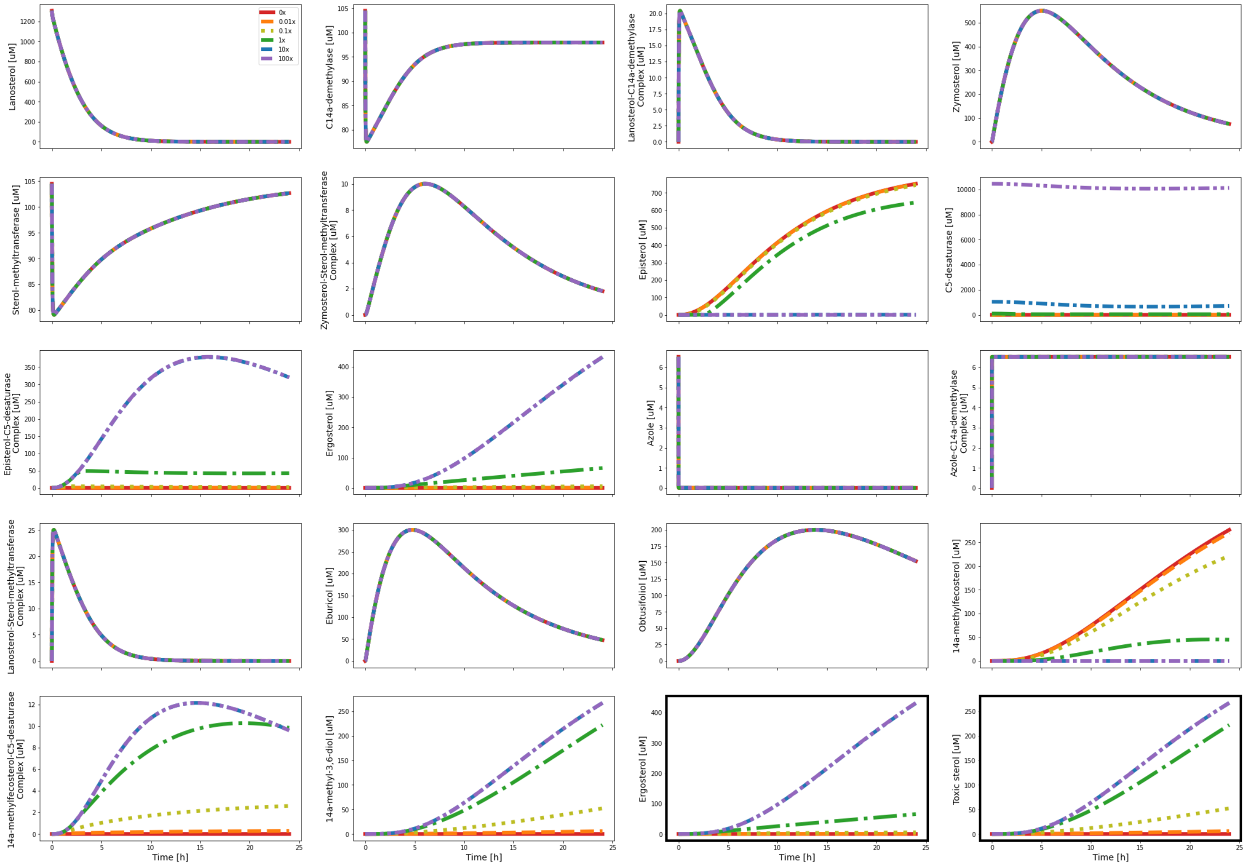


**Figure S7.** Plots generated by the simulation of variations in C5-desaturase concentration. The variations are as follows: 0x (red solid line), 0.01x (orange dashed line), 0.1x (yellow dotted line), 1x (green dashed dotted line), 10x (blue dense dashed dotted line), and 100x (purple dashed double dotted line). The x-axes denote the time in hours, while the y-axes represent the concentrations in µM. The last two subplots, emphasized in dark borders, reiterate the pertinent plots of ergosterol and the toxic sterol (14α-methyl-3,6-diol) for easier comparison.
